# Fourier shell correlation criteria for local resolution estimation

**DOI:** 10.1101/2020.03.01.972067

**Authors:** Alexis Rohou

## Abstract

The resolution of three-dimensional (3D) maps obtained by electron microscopy and reconstruction is rarely constant, owing to the flexibility or compositional variability of the macromolecules of interest. Accurately estimating the local resolution within 3D reconstruction volumes is desirable for analytical purposes, and may help improve the reconstruction process itself. One approach to estimating the local resolution is to compute Fourier shell correlations (FSC) between equivalent local regions of maps computed from half datasets: the estimated local resolution is that of the Fourier shell where the FSC drops below a pre-established threshold value. Existing thresholds can lead to significant over-estimates of the local resolution because they do not take into account the low number of independent voxels in low-resolution Fourier shells typically encountered when dealing with small sub-volumes. We derive a modified FSC threshold criterion, and a simple statistical test of whether the FSC significantly exceeds it, which aims to remedy this by taking into account the bias and variance of the FSC as an estimator of signal-to-noise ratio.

---

When computing or interpreting three-dimensional (3D) maps of macro-molecular structures obtained from electron microscopy images, the question often arises of what the resolution of the map, or of a part of the map, is. The most common way to estimate the resolution is somewhat indirect and involves the Fourier shell correlation (FSC) (Frank and Al-Ali, 1975; Harauz and Van Heel, 1986): the input image data are split and used to compute two 3D maps, which are compared with each other in a set of concentric shells in Fourier space. At each shell (corresponding to a narrow band of spatial frequencies), the correlation between the two “half-maps” is used as a proxy for the signal-to-noise ratio (SNR) of the reconstruction. The FSC, when plotted as a function of spatial frequency, begins at 1 at the origin and decays towards 0. A threshold value or curve is then used as a “resolution criterion”: the inverse of the spatial frequency at which the FSC first crosses the threshold is called the resolution.

Conducting this FSC experiment between two half-maps gives an estimated global resolution for the final map. To obtain estimates of the local resolution within regions of the map, small “sub volumes” can be extracted from each of the half-maps at equivalent real-space positions, and used to compute FSC curves. A similar test as described above is conducted to obtain an estimated resolution at each position of the map.

In this manuscript I revisit the use of “resolution criteria” originally developed for global resolution estimation in experiments to estimate local resolution.

## 1. Theory

### 1.1. The Fourier shell correlation

The Fourier shell correlation (FSC) between the Fourier transforms of two volumes, *F*_1_ and *F*_2_, is the sample correlation coefficient between Fourier components as a function of *s*, the Fourier shell index:

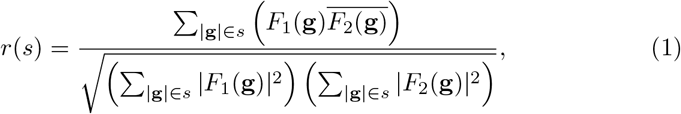

where **g** is a vector with spatial frequency |**g**|, and ̅ denotes complex conjugation. The FSC, *r*(*s*), is an estimate of the (unknown) population correlation *ρ*(*s*). In the following sections we will consider the statistical behavior of *r* within a single Fourier shell and therefore drop the *s* notation for clarity.

### 1.2. Correlation between half-dataset maps

The most common use of the FSC is to estimate the population correlation coefficient *ρ* between Fourier shells of two 3D reconstructions computed from two halves of a single dataset. The two “half maps” have been refined independently from each other, except at the lowest spatial frequencies. In some approaches the lowest spatial frequency at which the maps are refined independently is fixed (e.g., 40 Å by default in RELION; Scheres (2012)), but in others this parameter is adjusted during refinement (e.g. cisTEM; Grant et al. (2018)). In the next few sections, we will describe a number of relationships between the FSC, related quantities and estimates thereof, but these relationships only hold for those Fourier shells where maps are truly independent.

### 1.3. Correlation between the final map and the “ideal” map

It is useful first to consider *ρ*_ref_, the population correlation between a map computed from the full dataset (with Fourier components *F*_full_) and an ideal, error-free map (with Fourier components *F*_ref_). In reality such a map does not exist, but it can be simulated and serves as a useful thought experiment.

Rosenthal and Henderson (2003) showed that

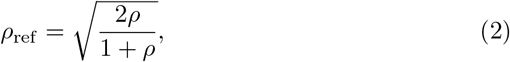

and proposed *ρ*_ref_ = 0.5 as a resolution threshold, yielding the widely-used threshold *ρ*_*t*_ = 0.143. To understand how to correctly utilize such a threshold under various conditions, it is helpful to consider the relationship between correlation and the signal-to-noise ratio.

### 1.4. Correlation and signal-to-noise ratio

Bershad and Rockmore (1974) showed that, under the conditions described above, where two versions of the same signal are corrupted by independent realizations of noise ^1^, the ensemble signal-to-noise ratio (SNR) and correlation are related by

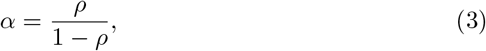

where *α* is the SNR in each of the shells (the SNR in the final reconstruction from the full dataset would be 2*α*).

Therefore the FSC threshold *ρ*_*t*_ = 0.143 may also be described as an SNR threshold *α*_*t*_ = 0.167. Using this as a resolution criterion is to ascertain at each shell whether *α* > *α*_*t*_. Of course *α* is not known, nor can it be calculated or measured directly; it can only be estimated. As we will describe below, there exist several estimators of *α*, with errors which may be significant and affect the outcome of the *α* > *α*_*t*_ test.

### 1.5. The FSC as a statistical estimator of the signal-to-noise ratio

Bershad and Rockmore (1974) showed that the sample correlation coefficient *r* can be used to construct a biased estimator of *α*, the signal-to-noise ratio in each of the half-maps being compared:

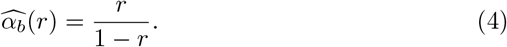

They also derived an unbiased version of this estimator, 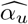, which is a function of *ρ*, the (unknown) population correlation coefficient, and *n*, the number of independent Fourier voxels within the shell:

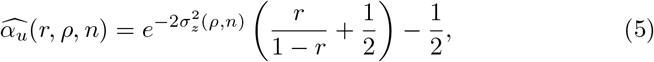

where

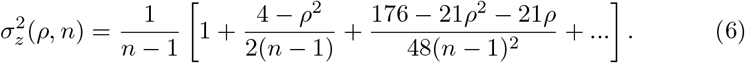

When using the FSC to estimate the global resolution of a 3D reconstruction, *n* is typically large (on the order of 10^2^), so that 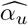 tends to 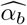, which justifies the use of the biased estimator (Equation 4) in much of the literature discussing resolution and SNR estimation in electron microscopy, beginning with Frank and Al-Ali (1975).

However when computing a local FSC over small sub-volumes (with sides on the order of 25 Å), and especially in low-resolution (> 15 Å) regions of maps, *n* can be very small (order of 10 voxels, see Table 1) in the relevant (low radius) Fourier shells. Under those conditions, 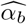 is significantly biased, i.e. it diverges from 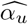 and over-estimates *α* on the order of 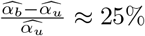.

**Table 1:**
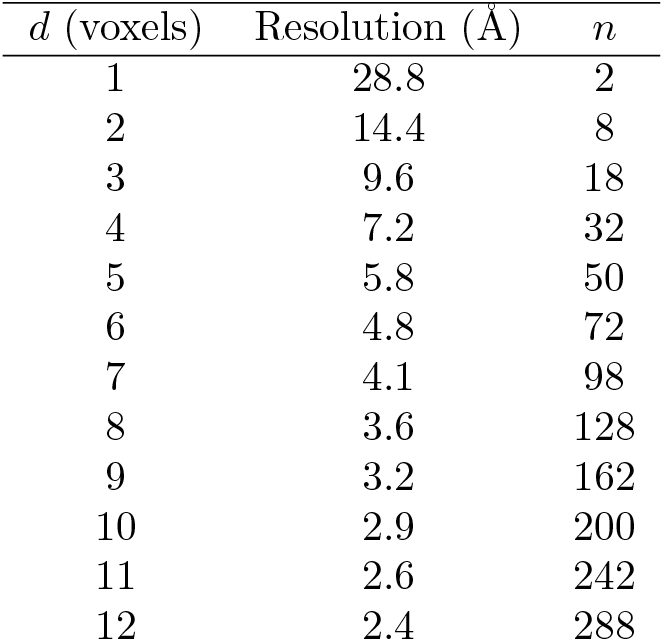
Resolution and number of independent voxels for the first few shells of a 24-pixel box with pixel size 1.2 Å. The number of voxels per shell is approximated as 2*πd*^2^Δ, where Δ = 1 is the thickness of a shell in voxels, *d* is the distance from the origin. When the FSC volumes are apodized in real space (as is common when measuring local resolution), the number of independent voxels is reduced. Here we assume that apodization causes about 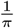 of the measurements to be independent, so that *n* = 2*d*^2^

| $d$ (voxels) | Resolution (Å) | $n$ |
| --- | --- | --- |
| 1 | 28.8 | 2 |
| 2 | 14.4 | 8 |
| 3 | 9.6 | 18 |
| 4 | 7.2 | 32 |
| 5 | 5.8 | 50 |
| 6 | 4.8 | 72 |
| 7 | 4.1 | 98 |
| 8 | 3.6 | 128 |
| 9 | 3.2 | 162 |
| 10 | 2.9 | 200 |
| 11 | 2.6 | 242 |
| 12 | 2.4 | 288 |

### 1.6. Constructing an FSC threshold based on the unbiased estimator

Rather than using the biased estimator (leading to *ρ*_*t*_ = 0.143), I propose to choose the FSC resolution threshold *ρ*_*t*_ so that

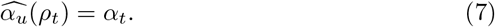

Substituting equation 5 and solving for *ρ*_*t*_ gives

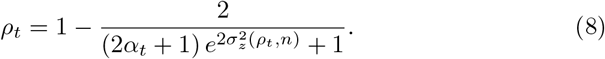

Further simplification is possible if one assumes that *n* is large, in which case 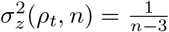:

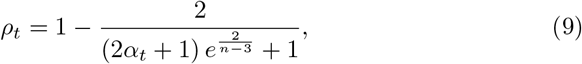

but this is not desirable since *σ*_*z*_ will then be mis-estimated in the low-resolution shells ^2^. Instead, equation 8 can be approximated numerically, as laid out in Appendix A. It is plotted in Figure 1, which suggests that the commonly-used fixed threshold *ρ*_*t*_ = 0.143 suffers from noticeable bias at resolutions of 5 Å or worse when used for local resolution estimation under the conditions described in Table 1.

**Figure 1:**
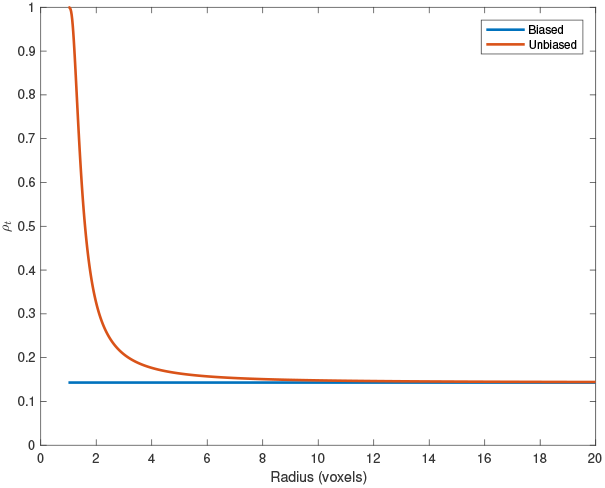
Threshold correlation values *ρ*_*t*_ derived using the biased and unbiased estimators for *α*_*t*_ = 0.167, equivalent to the familiar *ρ*_*t*_ = 0.143 threshold.

### 1.7. Significance tests

Having decided on a threshold for the FSC, one must construct a statistical test at each shell to decide whether the measured FSC, *r*, is significantly above the chosen threshold *ρ*_*t*_ so that one may be confident that *ρ* > *ρ*_*t*_. Equivalently, the test aims to ascertain whether the estimated SNR 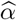 is significantly above the chosen threshold *α*_*t*_ so that one may be confident that *α* > *α*_*t*_.

In current usage, the last shell at which the measurement is above the threshold (*r* > *ρ*_*t*_) is used to determine the “resolution”. Implicitly, this test corresponds to a *p* value of approximately 0.5; to paraphrase, there is a 50% chance that, actually, the null hypothesis is true and that *ρ* < *ρ*_*t*_ at that shell.

It is possible to formulate more stringent statistical tests, for example requiring a 0.95 confidence level (i.e. *p* < 0.05) that *ρ* > *ρ*_*t*_, as was done by Penczek (2010), who derived the 95% confidence interval of an FSC curve based on the biased estimator 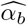 and the large-*n* approximation to 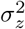.

Alternatively, we propose a test to ascertain whether 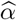 is siginificantly above *α*_*t*_.

### 1.8. A significance test based on 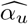

When doing a single FSC experiment to estimate the true SNR, the estimated SNR 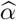 will be off relative to the true SNR *α*; the estimator will be in error: *r* ≠ *ρ* and therefore 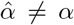. What is the probable magnitude of this error? Bershad and Rockmore (1974, Equation 14) derived the variance of their unbiased estimator:

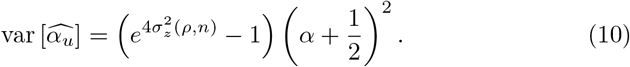

We propose setting the threshold *ρ*_*t*_ such that the estimated SNR 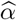 should be more than *k* standard deviations above the threshold SNR:

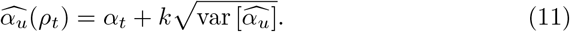

This yields

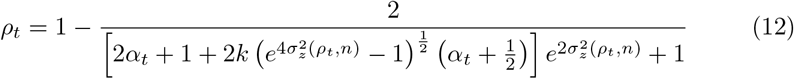

and its approximation, when *n* is large:

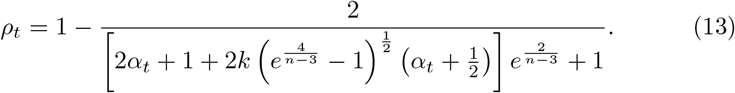

Experimentally measured FSC values (*r*) which are greater than *ρ*_*t*_ defined Equation 12 could then be taken to indicate with high certainty that the true SNR (*α*) in the reconstruction is actually greater than the desired threshold SNR (*α*_*t*_), even under conditions where the bias and error of the estimator are significant due to low *n*.

## 2. Results

### 2.1. Local resolution estimation programs

A new command-line program `local_resolution` was added to *cis*TEM (Grant et al., 2018). `local_resolution` re-implements the algorithm of Cardone et al. (2013) but with the option to use Equation 12 as a resolution criterion instead of fixed-value *ρ*_*t*_ criterion. Beyond this modification, several smaller changes were implemented relative to the original description (Cardone et al., 2013).

First, the input half-volumes are filtered so as to whiten their radial spectra, which improves the locality of resolution estimates (data not shown).

Second, we give the option to only estimate the local resolution every *m* voxels in every direction, resulting in a *m*^3^ time saving (this is also implemented in current versions of `blocres`), and to run the program in parallel, with each instance processing a slab of slices through the volume. A separate program `local_resolution_finalize` collates the results into a single output volume and interpolates (linearly) between voxels where the local resolution was estimated. Using typical settings (*m* = 2, pixel size = 1.3, box size = 20 pixels), a finalized local resolution map is obtained in X seconds using 4 Xeon processors (Intel).

Third, no zero-padding of subvolumes is used.

### 2.2. Behavior in low-resolution regions of maps

When using fixed thresholds, resolution estimates appear to be heavily biased against the very lowest frequencies. For example, in the case of a membrane-protein reconstruction (pixel size 1.2 Å, box size 24), I found that no voxels were estimated to have local resolutions lower than 11.3 Å when using *ρ*_*t*_ = 0.5. This was not the case when using equation 12 as a threshold (data not shown).

In 3D maps of membrane proteins, I have found that fixed thresholds in common use today (*ρ*_*t*_ = 0.5) typically estimate the resolution within detergent micelles to be ≈ 8-10 Å, which, if true, would suggest significant internal order, whereas using equation 12 typically gives estimates in the 15-30 Å range, likely corresponding to the envelope of the micelles, which may be presumed to be disordered internally.

## 3. Discussion

### 3.1. Which resolution threshold?

To my knowledge, van Heel and Schatz (2005) were the first to describe in detail how wrong the assumption can be that *n* is large and to conclude that fixed-value thresholds derived from the biased estimator 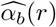 (Equation 5) should not be used. As I show above, the problem is especially acute when conducting local SNR estimates with small boxes; it would seem there are no good reasons to keep using fixed-FSC thresholds for local resolution estimation, other than programming simplicity and scientific inertia.

In that same paper, the authors also propose a new resolution threshold: the “half-bit criterion”, calibrated to denote the resolution at which each Fourier voxel in a shell still contributes at least 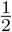 bit of information on average to the final map. In my view however, the two issues (of inaccurate SNR estimators on the one hand, and of resolution criteria on the other) are distinct and should be considered separately. The first issue may be corrected by forgoing fixed-FSC resolution thresholds and using instead the appropriate unbiased estimator as outlined above.

This can be done whether following Crowther, Rosenthal and Henderson (Rosenthal and Henderson, 2003, Appendix) in choosing *α*_*t*_ = 0.167 as an SNR threshold, or following van Heel & Schatz in choosing *α*_*t*_ = 0.2071. In fact, using *α*_*t*_ = 0.167 and *k* = 2 yields a threshold curve numerically similar to the half-bit curve (Figure 3), demonstrating that the choice of SNR or (information content) threshold is conceptually orthogonal to the choice of SNR estimator (biased versus unbiased) and of statistical testing framework.

### 3.2. Lack of confidence measures in the literature

To my knowledge, all single-particle cryoEM image processing software packages in use today perform the *α* > *α*_*t*_ test by checking 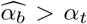 at every shell without considering the confidence with which one can reject the null hypothesis that in fact *α* ≤ *α*_*t*_. This is despite a method for doing so having been proposed some time ago by Penczek (2010, Figure 3.2). To caricature slightly, one might say that all (global and local) resolution estimates so far have a *p* value of *p* ≈ 0.5, or a *k* of 0 in the nomenclature introduced here. Recent proposals (Ludtke, Sorzano, and others, communications via listservers) to include, e.g., error bars in FSC figures could remedy this somewhat, but would leave unsolved the problem of bias in SNR estimators.

**Figure 2:**
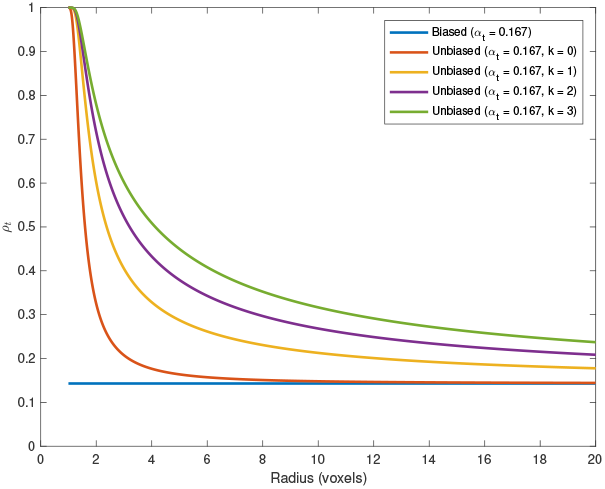
Correlation thresholds with a target SNR of *α*_*t*_ = 0.167. Using the biased SNR estimator (Equation 4) leads to the familiar *ρ*_*t*_ = 0.143 threshold, which is significantly biased in the lowest shells and is very permissive (*p*-value ≈ 0.5).

**Figure 3:**
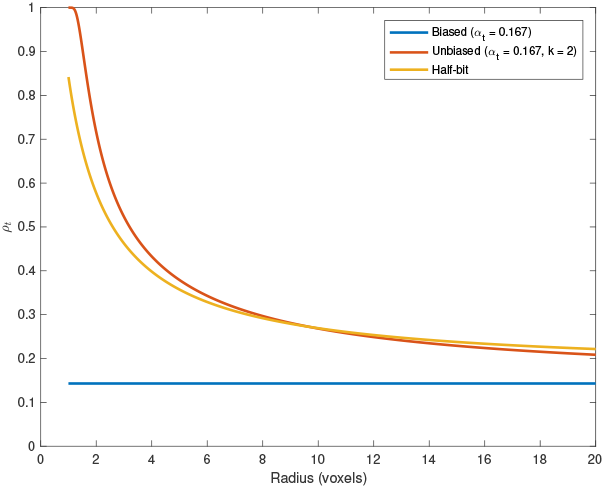
A correlation threshold using the unbiased estimator, a target *α*_*t*_ = 0.167 (akin to *ρ*_*t*_ = 0.143) and *k* = 2 is numerically similar to the “half-bit” criterion of van Heel and Schatz (2005)

### 3.3. Using improved estimators and significance tests for global resolution estimation

van Heel and Schatz (2005) pointed out that, even in the context of experiments to estimate the global resolution of a map, symmetry and other effects can cause *n* (the number of independent Fourier voxels in a shell) to be small and FSC measurements to behave in ways that are not reflected by the biased SNR estimator (Equation 4) or the fixed FSC threshold criteria derived from it.

While most global resolution claims today in the field of single-particle cryo-EM likely do not suffer significantly from this problem (because the claims are based on FSC measurements in shells where *n* is large), if a non-fixed FSC threshold based on unbiased estimates of SNR, preferably combined with a significance test, were to be agreed upon and adopted by software packages in the field, the risk of erroneous resolution claims might be expected to decrease, particularly in the context of “edge cases” for example involving small (possibly masked) objects reconstructed in large boxes, and/or highly-symmetrical maps.

### 3.4. When is n too low for resolution estimation, even when using unbiased estimators?

The significance test described above (*r* > *ρ*_*t*_ as defined in Equation 12) is not numerically stable when *n* is very low (e.g., for resolutions worse than 15 Å under the conditions used for Table 1), because *ρ*_*t*_ and *r* both approach 1 and *n* approaches 0. It would be desirable to derive the limit (smallest *n*, largest *ρ*_*t*_, or both) below which the local resolution should not be tested using current parameters. For regions of maps where this is the case, larger box sizes could be used so that *n* would be larger for the relevant local resolutions.

### 3.5. Other open questions

It is unclear what should motivate the choice of *α*_*t*_. Crowther, Rosenthal and Henderson (Rosenthal and Henderson, 2003, Appendix) proposed a criterion based on considerations from the field of macromolecular x-ray crystallography, whereas van Heel and Schatz (2005) chose *α*_*t*_ based on the average estimated information content of Fourier components. Either choice of *α*_*t*_ is compatible with the framework described in this manuscript, but neither specifically addresses the relationship between spectral SNR and real-space resolvability of features following Fourier synthesis.

## 4. Acknowledgments

Thanks to Tim Grant and Evan Green for testing early implementations of the algorithm, to Tim Grant for the suggestion to whiten the input half-maps, and to Sara L. Campbell for reviewing an earlier draft of the manuscript.

## Appendix A. Numerical approximation of 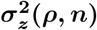

To be written.

## Footnotes

1 and therefore 0 < *ρ* ≤ 1, and *α* > 0

2 It turns out that using the simplification overestimates the variance of *z*, the Fisher transform of *r*. For example, when *n* = 10 and *ρ* = 0.9, the 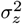 is approximated as 0.143 rather than the more accurate 0.1348, an overestimate of 1.06×.

